# A modular neural circuit for computing the motion of objects

**DOI:** 10.64898/2026.06.07.730718

**Authors:** Ethan Trepka, Catherine Yue, Ruobing Xia, Shude Zhu, Sharif Saleki, Danielle Abreu Lopes, Stephen Niño Cital, Tirin Moore

**Affiliations:** Neuroscience Interdepartmental Program, Stanford University, Stanford, CA, USA; Howard Hughes Medical Institute, Stanford University, Stanford, CA, USA; Department of Neurobiology, Stanford University, Stanford, CA, USA

**Keywords:** Motion Processing, Neuronal Dynamics, Visual Cortex

## Abstract

Computing the direction of a moving object requires integrating motion signals from its component edges into coherent patterns. The representation of pattern motion in visual cortex has been extensively studied, yet its underlying neural circuitry remains unknown. Using high-density recordings in macaque area MT, we show that selectivity to pattern motion emerges from a functionally and anatomically distinct cortical circuit. We demonstrate that neurons specialized for encoding component and pattern motion comprise distinct cell types arranged in a hierarchical circuit in which pattern neurons integrate inputs from a range of component neurons. Furthermore, we show that component and pattern neurons are spatially segregated into modules arranged systematically across cortical columns encoding direction of motion. This architecture and circuit align with a classic solution to the problem of computing the motion of objects.

## Introduction

Visual perception emerges from a series of transformations across structures in the brain. A classic exemplar in the mammalian visual system is the evolution of neuronal response properties from the lateral geniculate nucleus to primary visual cortex (V1), whereby circular center-surround receptive fields in the former give rise to orientation-selective simple and complex cells in the latter (Hubel and Wiesel 1962; Martinez and Alonso 2001; Alonso and Martinez 1998; Hubel and Wiesel 1959). These circuits provide the foundation for more complex visual computations, such as extracting the motion of objects.

In motion perception, computing the direction of a moving object requires integrating motion signals from its component edges into coherent patterns (Marr and Ullman 1981; Adelson and Movshon 1982). Across mammalian species, neurons at early stages in the visual system are most sensitive to the motion of component edges and at later stages become sensitive to the motion of patterns (Solomon et al. 2011; Lempel and Nielsen 2021; Juavinett and Callaway 2015; Rodman and Albright 1989; Movshon et al. 1985; Huk and Heeger 2002; Castelo-Branco et al. 2002). This transformation from component to pattern representations has been particularly well studied in the macaque visual system. Whereas neurons in macaque V1 respond principally to the motion of the components in a complex pattern (Movshon et al. 1985; Movshon and Newsome 1996), a substantial proportion of neurons downstream in area MT respond to the motion of the pattern itself (Rodman and Albright 1989; Movshon et al. 1985). This transformation is thought to play an essential role in computing the motion of whole objects (Born and Bradley 2005; Movshon et al. 1985; Rodman and Albright 1989). Yet, the circuitry underlying this transformation remains unknown.

Questions about the emergence of pattern representations in area MT have persisted since their discovery (Wang and Movshon 2016; Rust et al. 2006; McDonald et al. 2014; Wang 1997; Movshon et al. 1985; Simoncelli and Heeger 1998; Quaia et al. 2022). For example, pattern neurons could reflect a continuum of selectivity from component to pattern motion. Alternatively, pattern neurons could represent a distinct class of neurons that emerge from a hierarchical transformation of component to pattern representations within MT. Resolving such questions is necessary for understanding the computation of object motion and elucidating the nature of hierarchical transformations in the visual system. But doing so would require more comprehensive measurements from large distributed populations of neurons, which was not possible in past studies.

In recent years, high-channel-count electrophysiological recording devices such as Neuropixels probes have enabled simultaneous recordings from large, local populations of neurons across the brain (Trautmann et al. 2025; Jun et al. 2017). High-density recordings provide measurements of the activity of individual neurons in the context of a broader circuit. Thus far, they have been leveraged to characterize the functional architecture and circuitry of brain structures across species (Leonard et al. 2024; Siegle et al. 2021; Namima et al. 2025; Carr et al. 2026). Such recordings are particularly well-suited for use in the macaque visual system, where a large fraction of visual areas, including area MT, are located within deep and convoluted neocortex. Thus, we leveraged high-density recordings to elucidate how visual cortical circuits compute the motion of objects.

## Results

### High-density recordings reveal modular organization of area MT

We used the recently developed Neuropixels NHP 1.0 probe to measure the visual properties of 2,096 direction-selective neurons in area MT across 17 sessions in two macaque monkeys (A & H) (**Methods**) (**Fig. S1**). Our recordings utilized 384 channels near the tip of the Neuropixels probe spaced ∼20 µm vertically and ∼100 µm horizontally along the shank (**Fig. 1a**), providing high-density sampling of the tuning properties of MT neurons. By examining the direction of motion tuning of neurons along the length of the probe, we observed that nearby neurons tend to have similar preferred directions as predicted by MT’s established columnar organization (Albright et al. 1984; Diogo et al. 2003; DeAngelis and Newsome 1999). The systematic variation in preferred direction across the length of the electrode was apparent in individual sessions (**Fig. 1b; Fig. S2**). Notably, these recordings also revealed occasional ∼180° reversals, or so-called ‘fractures’ (Hubel et al. 1978; Diogo et al. 2003; Albright et al. 1984), in preferred motion direction between neighboring recording sites. We utilized an automated approach to detect and label these reversals across recordings (**Methods**) (48 total reversals, 3.43 reversals/session).

**Figure 1.**
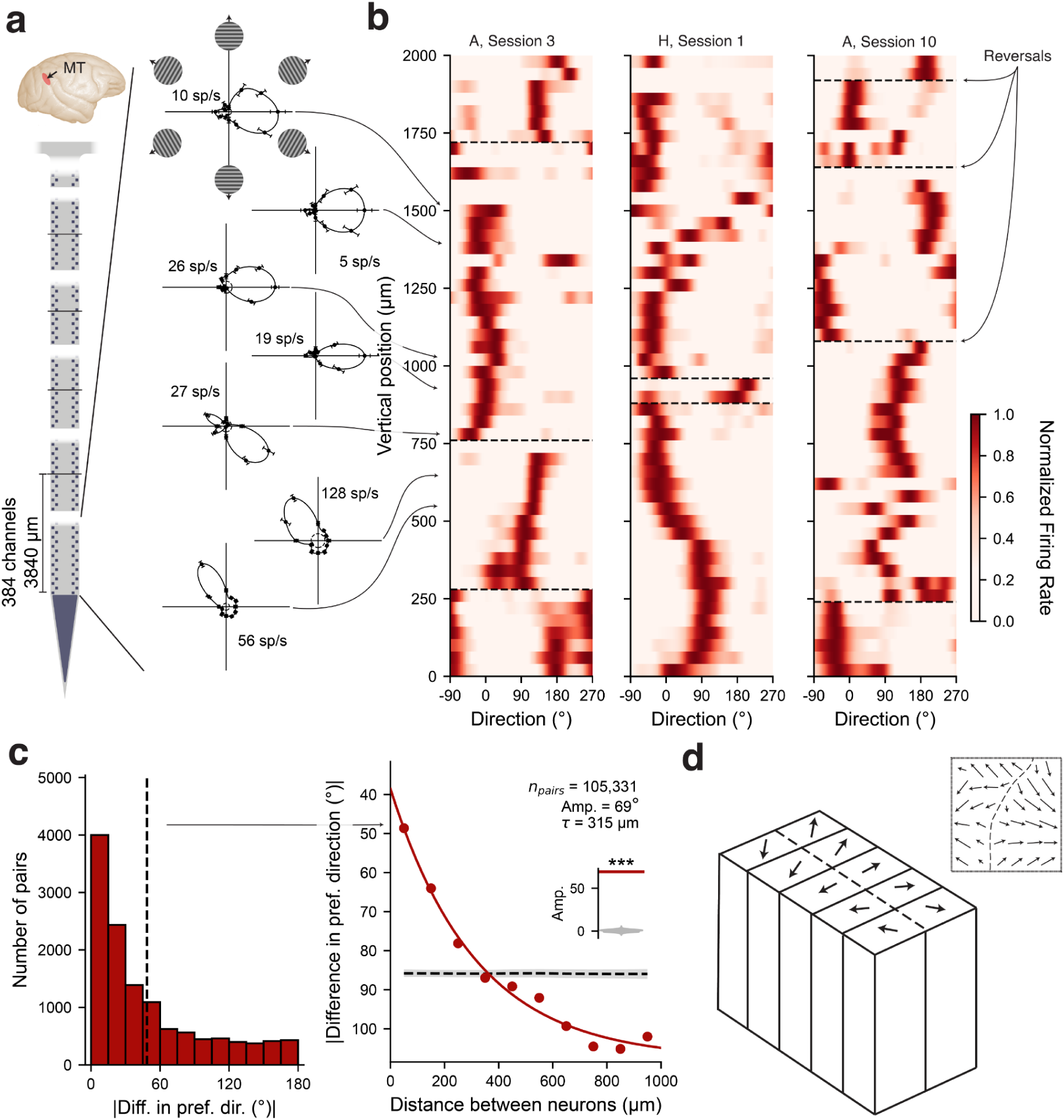
Modular organization of motion direction tuning in area MT. **(a)** Top left, the location of the middle temporal visual area (MT) (red shaded) in the superior temporal sulcus (projected onto the cortical surface for visualization). Bottom left, schematic of Neuropixels NHP probe, with the 384 channels near the tip of the probe used for recordings indicated. Right, example tuning curves for six neurons to grating drift direction, ordered by their position relative to the probe tip. Dashed circle denotes the baseline firing rate, text denotes the neuron’s average firing rate to its preferred grating direction, and error bars denote SEM. **(b)** Grating direction tuning curves for all neurons at positions relative to the probe tip in 40 μm bins, in three example sessions. Tuning curves for each neuron are normalized between 0 and 1 by subtracting the neuron’s minimum firing rate across direction conditions and dividing by its maximum.

To quantify the modularity and spatial scale of direction of motion tuning, we examined the difference in preferred direction between pairs of neurons as a function of the distance between neurons (**Fig. 1c**) (105,331 total pairs, 6,196 pairs/session). The difference in preferred direction decayed exponentially, with a spatial scale (𝜏) of 315 µm (R^2^=0.98). The amplitude of the exponential fit (69°) was significantly greater than that of a label-shuffled null model (permutation test, p < 0.001). Notably, the magnitude of the distance relationship we observed (e.g., difference > 100° at 1 mm) is consistent with previous data from tangential penetrations of MT (DeAngelis and Newsome 1999), suggesting that our penetrations were largely tangential to the cortical surface. Our data are thus consistent with the classic model of MT’s functional organization in which smooth changes in preferred direction across the cortical surface coexist with abrupt reversals (Albright et al. 1984; Diogo et al. 2003) (**Fig. 1d**). These results also demonstrate that high-density recordings can reveal the modularity of visual properties within cortex in single recordings.

Dashed black lines indicate 180° reversals. **(c)** Left, distribution of the difference in preferred direction of all pairs of neurons with a distance less than 100 μm, aggregated across all sessions. The dashed black line indicates the mean difference in preferred direction. Right, distance between neurons vs mean difference in preferred direction, aggregated across all sessions. Note that larger differences in preferred direction correspond to lower values on the y-axis. The black dashed line and gray shaded region denote the baseline and 95% confidence interval from a label-shuffled null model. The red line denotes an exponential fit. Amplitude and spatial scale (𝜏) of the fit are denoted, and the inset illustrates observed amplitude (red) vs the distribution of amplitudes of the shuffled null (gray) (*** indicate p < 0.001, permutation test). **(d)** Schematic representation of the hypercolumnar organization of direction of motion selectivity in area MT (Albright et al. 1984). Direction preference changes smoothly tangential to the cortical surface, with occasional 180° reversals (dashed line). Inset illustrates additional complexity present in the organization of direction of motion selectivity in area MT, based on data from (Diogo et al. 2003).

### Measurement of component and pattern motion selectivity with high-density recordings

We next characterized component and pattern motion selectivity by measuring neuronal responses to drifting gratings and to plaid stimuli constructed by superimposing two drifting gratings with drift directions offset by 120°, following previously established methods (Smith et al. 2005) (**Methods**) (**Fig. 2a**). In a plaid stimulus, each component motion signal constrains, but does not uniquely determine, the direction of the overall pattern. The pattern motion direction of the plaid can be inferred by integrating the two component signals (Marr and Ullman 1981; Adelson and Movshon 1982).

**Figure 2.**
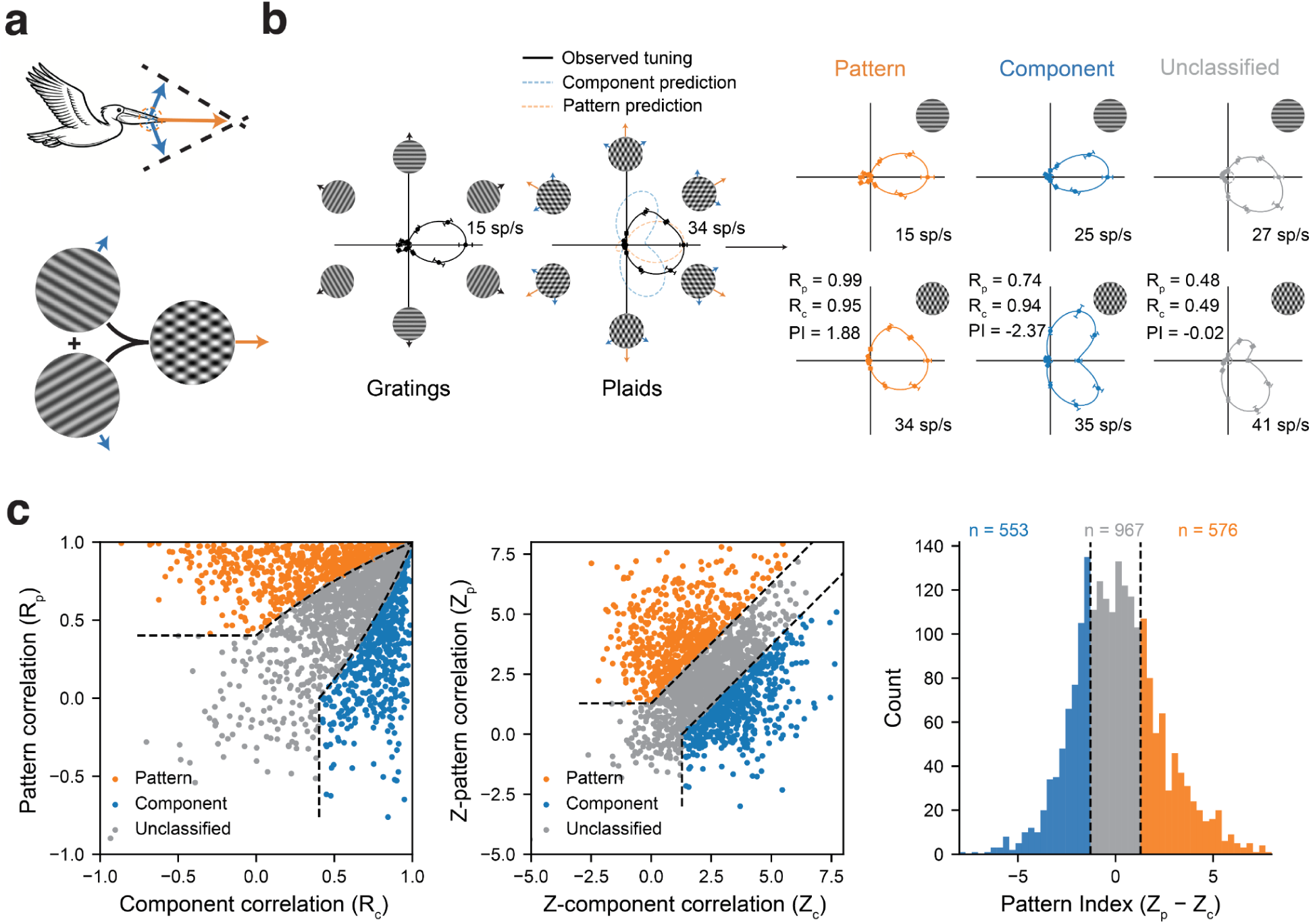
Recordings from large populations of neurons specialized for encoding component and pattern motion. **(a)** Top, schematic of the computation of object motion from edge motion. An object’s motion direction (orange vector) is given by the intersection of constraints (dashed) defined by motion orthogonal to its component edges (blue vectors). Bottom, drifting grating and plaid stimuli used to probe neurons. A plaid stimulus is constructed by superimposing two drifting gratings with drift directions offset by 120°. **(b)** Left, measured tuning curves for grating and plaid motion direction for an example neuron. Predicted plaid tuning curves for a component or pattern neuron are overlaid. Dashed circle denotes the baseline firing rate, text denotes the firing rate in the preferred condition, and error bars denote SEM. Right, measured tuning curves for grating (top) and plaid (bottom) motion direction for example pattern (orange), component (blue), and unclassified (gray) neurons. The component and pattern partial correlations (R_c_, R_p_) and the neuron’s pattern index (PI) are indicated. **(c)** Classification of pattern and component neurons based on the raw (left) and z-transformed (middle) component and pattern partial correlations (R_c_, R_p_ and Z_c_, Z_p_, respectively), and the pattern index (Z_p_ - Z_c_) (right) across neurons in all sessions (n = 2,096). The dashed black lines indicate the classification boundaries separating pattern, unclassified, and component neurons.

In our large-scale dataset, we observed neurons that could be classified as pattern, component, or unclassified based on how well their response to plaids fit the component or pattern predictions (**Methods**) (**Fig. 2b**), similar to previous single-electrode studies (Smith et al. 2005; Wang and Movshon 2016; Movshon et al. 1985; Rodman and Albright 1989; Pack et al. 2001; Lempel and Nielsen 2021; Juavinett and Callaway 2015). The predicted tuning curve of a pattern neuron to plaids contains a single peak aligned to its preferred grating direction, whereas the predicted tuning curve of a component neuron contains two response peaks corresponding to the two plaids containing its preferred grating direction. The ‘pattern index’ for each neuron summarizes its selectivity, with values above 1.28 corresponding to pattern neurons (Smith et al. 2005; Wang and Movshon 2016). Our recordings identified 553 component neurons and 576 pattern neurons across recording sessions (**Fig. 2c**). As observed in previous studies, pattern selectivity appeared continuously distributed across the population of single neurons (Wang and Movshon 2016; Smith et al. 2005). Importantly, substantial numbers of component and pattern neurons were identified in individual recording sessions (mean: 33 component and 34 pattern neurons per session).

### Modular organization of pattern and component neurons in area MT

We next asked whether pattern and component neurons are randomly intermixed or spatially organized within MT. When examining tuning to plaid direction along the length of the electrode in single sessions, we found that the two-peaked tuning curves (‘component neurons’) and single-peaked tuning curves (‘pattern neurons’) tended to fall into separate clusters (**Fig. 3a**). We visualized this effect by plotting the pattern index of all neurons along the length of the probe at each electrode contact where a neuron was detected. This revealed that the clustering of component and pattern neurons was consistent across recording sessions (**Fig. 3b; Fig. S3**).

**Figure 3.**
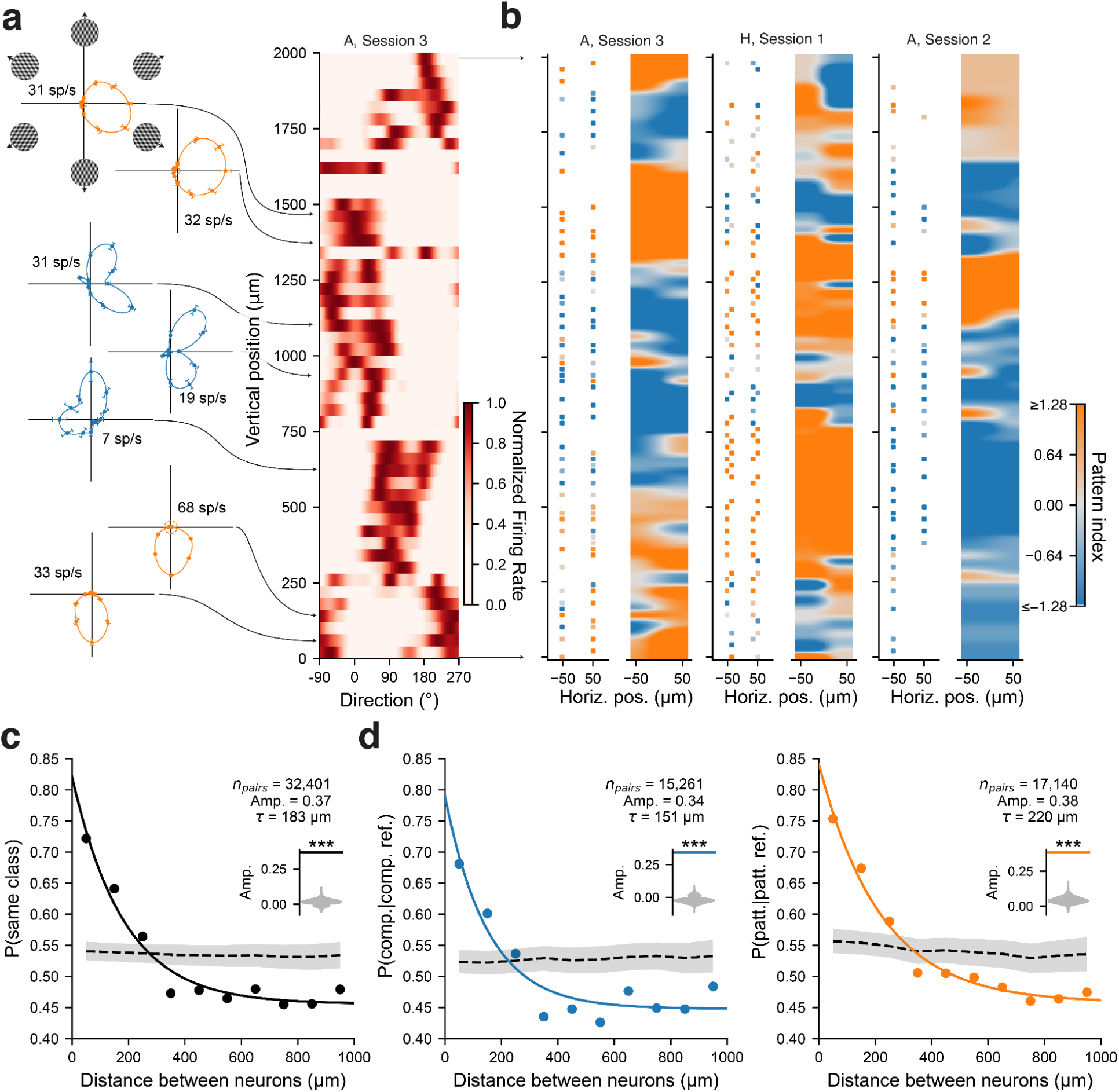
Modular organization of component and pattern neurons. **(a)** Left, example tuning curves for six neurons to plaid drift direction, ordered by their position relative to the probe tip and colored based on class (pattern orange, component blue). Dashed circle denotes the pre-stimulus baseline firing rate, text denotes the neuron’s firing rate in its preferred plaid condition, and error bars denote SEM. Right, plaid direction tuning for all neurons at positions relative to the probe tip in 40 μm bins in an example session. **(b)** Pattern index for all neurons at positions relative to the probe tip in three example sessions. Left panels, the Neuropixels probe contacts are colored based on the pattern index of the neuron detected on that contact. Contacts where a neuron was not detected are omitted. Right, interpolated pattern index vs electrode map. **(c)** Distance between pairs of neurons vs the probability that both neurons share the same class identity: (P(same class) = (#Patt.-Patt. + #Comp.-Comp.) / (#Patt.-Patt. + #Comp.-Comp. + #Patt.-Comp.)), aggregated across all sessions. The black dashed line and gray shaded region denote the baseline and 95% confidence interval from a label-shuffled null model. The black line is an exponential fit, and the amplitude and spatial scale (𝜏) of the fit are denoted. The inset illustrates measured amplitude (black) vs the distribution of amplitudes of the shuffled null (gray) (*** indicate p < 0.001, permutation test). **(d)** Left, distance between pairs of neurons vs the probability of the target neuron being a component neuron given that the reference neuron is a component neuron (P(Comp.|Comp. Ref.) = (#Comp.-Comp.) / (#Comp.-Comp. + #Patt.-Comp./2)). Right, similar but for a pattern neuron. Conventions in (d) are similar to (c).

To quantify this modularity and its spatial scale, we examined the probability that two neurons shared the same class (both pattern or both component) as a function of the distance between neuronal pairs. Remarkably, nearby pairs (∼100 µm apart) were much more likely ( ∼75%) to share the same class than distant pairs (∼1000 µm apart; ∼45%) (**Fig. 3c**). The probability of pairs sharing the same class decayed exponentially as a function of distance (R^2^=0.96), with a spatial scale (𝜏) of 183 µm, and the amplitude of the exponential fit (0.37) was significantly greater than that of the label-shuffled null model (permutation test, p < 0.001). The probability of neuronal pairs sharing the same class provides a summary metric for modularity that combines information about both pattern and component clusters. Nonetheless, this modularity could be due to modularity in only one of the two neuron classes. Thus, we examined the modularity of component and pattern neurons separately. We found that both classes of neurons were clustered into modules. Component clusters exhibited a spatial scale (𝜏) of 151 µm (R^2^=0.89) while pattern clusters exhibited a spatial scale (𝜏) of 220 µm (R^2^=0.98) (**Fig. 3d**). The amplitude of the exponential fit for both component (0.34) and pattern (0.38) clusters was significantly greater than that of a label-shuffled null model (permutation test, p < 0.001). The observed spatial modularity of pattern and component neurons provides the first evidence that these classes represent anatomically distinct populations of neurons.

### Distinct spike waveforms among pattern and component populations

Next, we asked whether spike waveforms differed between pattern and component neurons, which could reflect differences in underlying morphological cell types (Henze et al. 2000; Carr et al. 2026; Constantinidis and Goldman-Rakic 2002; Mitchell et al. 2007). Compared to single-or low-channel count electrodes, high-density recordings provide improved measurements of the heterogeneity of waveforms in large local populations, and thus are uniquely suited to address this question (Zhu et al. 2024; Carr et al. 2026; Paulk et al. 2022; Jun et al. 2017; Leonard et al. 2024) (**Fig. 4a**). First, we tested whether pattern and component neurons could be classified from their extracellular waveforms (**Fig. 4b**). To capture potential nonlinear relationships between waveform shape and functional class, we used a support vector machine with a radial basis kernel (**Methods**). This classifier could distinguish pattern and component cells solely from their waveforms (62% class-balanced accuracy; 50%, label-shuffled mean, p<0.01, permutation test) (**Fig. 4c**). To better understand what waveform features distinguish pattern and component neurons, we split extracellular waveforms into fast-spiking (FS; putative inhibitory) and regular-spiking (RS; putative excitatory) categories based on their trough-to-peak duration, and retrained the classifier on each subset of waveforms (**Fig. 4d**) (**Methods**) (Mitchell et al. 2007). The binary classifier trained on the FS subset of waveforms achieved an accuracy of 64% (**Fig. 4e**) and the classifier trained on the RS subset achieved an accuracy of 61% (**Fig. 4f**). We then looked for differences between pattern and component neurons within the FS and RS waveform subsets. This revealed that the peak amplitudes were significantly larger for pattern neurons than component neurons (med. diff.=0.03, p<0.001, Mann-Whitney U-test), and this difference was much more pronounced among FS neurons (med. diff.=0.09, p<0.001, Mann-Whitney U-test) than RS neurons (med. diff.=0.02, p<0.05, Mann-Whitney U-test) (**Fig. 4g**). Although the mapping between the peak amplitude of the waveform and cell type is not fully understood, variations in this region of the waveform are generally attributed to differences in morphology or active conductance distribution (Gold et al. 2006; Henze et al. 2000). The observed differences in spike waveforms between pattern and component neurons provide a second piece of evidence that these classes represent distinct populations of neurons.

**Figure 4.**
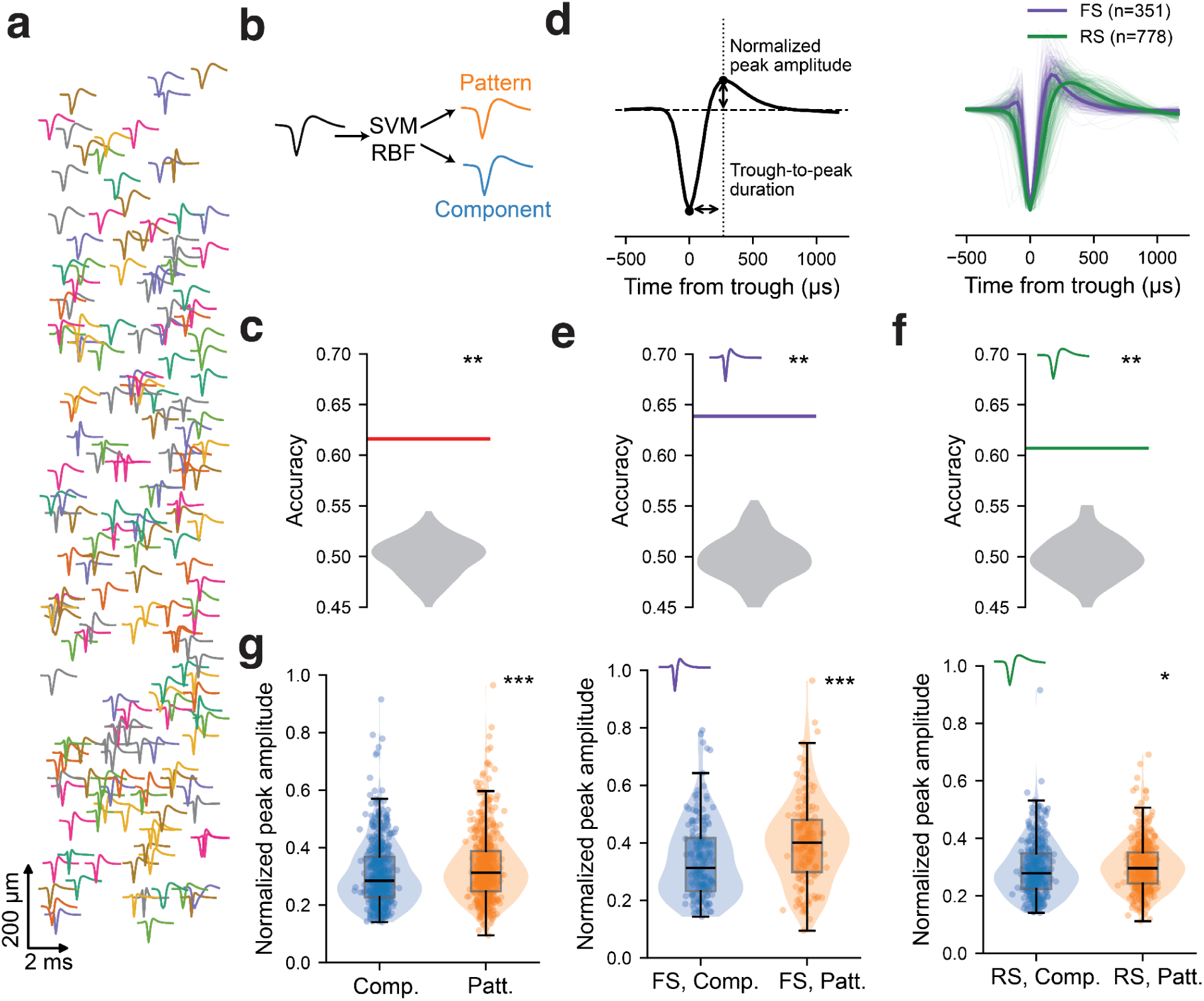
Component and pattern neuronal populations are composed of distinct spike waveforms. **(a)** Extracellular waveform templates for all recorded neurons in an example session, plotted at their position relative to the Neuropixels probe tip (y). The x-position is randomly jittered for visualization. **(b)** Schematic of classification task. Given an unlabeled extracellular waveform, a binary classifier (SVM RBF, support vector machine with a radial basis function kernel) assigns it to either the pattern or component category. **(c)** Accuracy of the binary classifier (solid red line) relative to a null distribution obtained from shuffling class labels and retraining the classifier (** indicate p < 0.01, permutation test). **(d)** Left, an example extracellular waveform and waveform features extracted from it, including trough-to-peak (TTP) duration and normalized peak amplitude. Right, mean fast-spiking (FS; TTP <= 200 µs) and regular-spiking (RS; TTP > 200 µs) waveforms, overlaid on example traces. **(e-f)** Accuracy of the binary classifier trained separately on the subset of FS (e) and RS (f) neurons. Conventions follow (c). **(g)** Normalized peak waveform amplitude for all (left), FS (middle), and RS (right) component and pattern neurons. Boxplots indicate first quartile, median, and third quartile, and scatter points correspond to individual neurons (* and *** indicate p < 0.05 and p < 0.001, respectively, Mann-Whitney U-test).

### Functional connections from component to pattern neurons

Given the evidence of pattern and component modularity and differences in spike waveforms, we next asked whether these populations exhibit different patterns of functional connections. Temporally precise correlations in spiking activity (e.g., cross-correlation) have long provided a means of assessing connections among neurons in the mammalian visual system (Usrey et al. 1998; Ts’o et al. 1986; Toyama et al. 1981; Siegle et al. 2021; Schwarz and Bolz 1991; Reid and Alonso 1995; Baker and Bair 2012; Alonso and Martinez 1998), e.g., by providing evidence of the circuit-level hierarchy from simple to complex cells in V1 (Alonso and Martinez 1998). The use of high-density recordings substantially increases the yield of functionally connected pairs identifiable in single recordings, thereby facilitating the characterization of circuit-level interactions (Siegle et al. 2021; Panichello et al. 2024; Carr et al. 2026). Thus, we leveraged our high-density recordings to examine the circuitry underlying the emergence of pattern selectivity in MT.

We computed cross-correlations between the activity of simultaneously recorded neurons within each session. We used activity in the late visual period (100-250 ms after stimulus onset) to ensure the analysis was independent of the visual transient and neuronal response latency. We defined functional connections between pairs of neurons by applying a significance test to each cross-correlation histogram, yielding a directed graph of functional connectivity within each session (**Fig. 5a**). A pair of neurons was classified as functionally connected if activity in one neuron (source) reliably led (1-5 ms) activity in another (target) and the peak significantly exceeded the baseline (**Methods**) (English et al. 2017).

**Figure 5.**
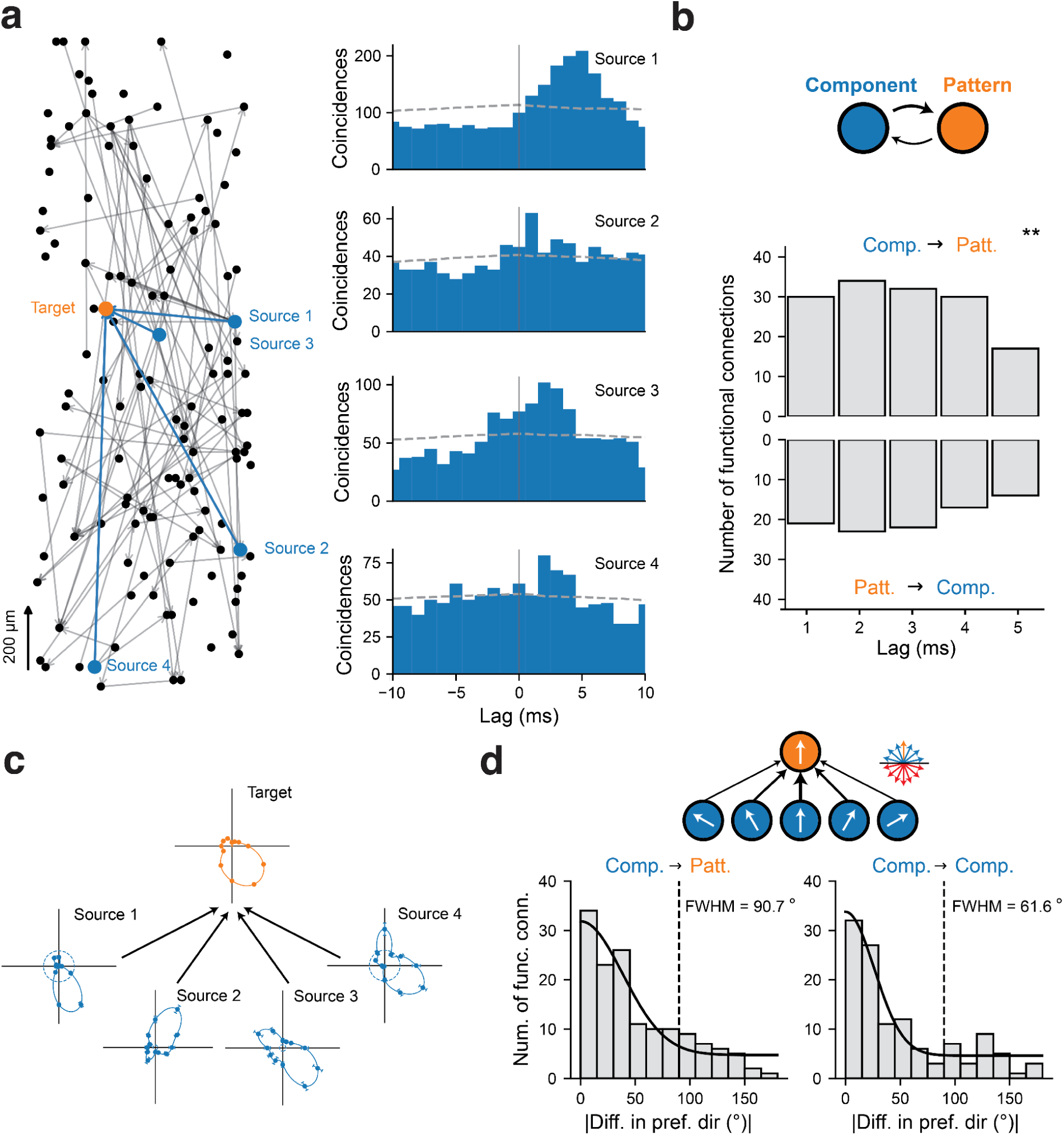
Functional connections between component and pattern neurons. **(a)** Left, directed graph of all functional connections in an example session. Neurons are plotted at their position relative to the Neuropixels probe tip (y) and their x-position is randomly jittered for visualization. The edges between a target pattern neuron and four component source neurons that it received functional connections from (component-> pattern functional connections) are highlighted in blue. Right, the cross-correlation histograms (CCH) for the four source component neurons and the target pattern neuron. A peak in the CCH at positive time lags corresponds to the source neuron firing prior to the target neuron. The gray dashed line denotes the slow baseline CCH obtained by convolving the CCH with a hollowed Gaussian. **(b)** Component neurons tend to lead pattern neurons. The distribution of lags for component-to-pattern and pattern-to-component functional connections is illustrated across all sessions. (** indicate p < 0.01, binomial test of number of component-to-pattern vs pattern-to-component connections) **(c)** Grating direction tuning curves for the four component source neurons and their target pattern neuron from (a). Tuning curve conventions follow Fig. 3a. **(d)** Histogram of difference in preferred grating direction between source and target neurons for component-to-pattern and component-to-component functional connections. Black solid line illustrates Gaussian fit and text denotes connectivity bandwidth (full-width at half-maximum, FWHM of the fitted Gaussian). Black dashed line denotes 90° difference in preferred grating direction. Schematic illustrates a pattern neuron that receives input from component neurons with preferred directions within 90° of its preferred pattern direction. Schematic inset denotes the set of component motion vectors consistent (blue) and inconsistent (red) with a given pattern motion direction (orange) under the intersections of constraints model.

In total, we identified 246 functionally connected component-pattern pairs out of 18,860 total possible component-pattern pairs (1.3%). Among these, we observed both component-to-pattern and pattern-to-component connections. Notably, there were significantly more component-to-pattern than pattern-to-component pairs (**Fig. 5b**) (n_component-to-pattern_=144, n_pattern-to-component_=102, ∼1.4:1 ratio, p < 0.01, binomial test). A similar bias was observed when activity in the early visual period was included (0-250 ms) (n_component-to-pattern_=260, n_pattern-to-component_=119, ∼2.2:1 ratio, p < 0.001, binomial test) (**Fig. S4**). These results indicate that the direction of functional connections between component and pattern neurons was biased, with component neurons tending to lead pattern neurons. The magnitude of the observed connection asymmetry is comparable to that of simple-to-complex cell connections in V1 (Trepka et al. 2022) and neurons in hierarchically adjacent areas of mouse visual cortex (Siegle et al. 2021). The observed differences in functional connectivity between pattern and component neurons provide a third piece of evidence that these classes represent distinct populations of neurons.

Computational models of pattern neurons predict that they integrate inputs from a broad range of component neurons with preferred directions within 90° of their preferred pattern direction (Rust et al. 2006; Simoncelli and Heeger 1998). Indeed, we observed pattern neurons that received functional input from component neurons with a broad range of preferred directions (**Fig. 5c**). To test this prediction across the full population, we examined the relationship between the preferred directions of the source and target neurons among the component-to-pattern pairs. Specifically, we quantified the breadth of functional inputs to pattern neurons by examining the distribution of difference in preferred direction among these pairs (**Fig. 5d**). A Gaussian fit to this distribution yielded a connectivity bandwidth of 91° (SE=13.17, R^2^=0.90). By comparison, for the set of component-to-component pairs (n=119), the connectivity bandwidth was 62° (SE=7.91, R^2^=0.92). Consistent with the connectivity results, we also found that the tuning bandwidth to grating stimuli was significantly larger for pattern neurons (median bandwidth=96°) than for component neurons (median bandwidth=66°) (p < 0.001, Mann-Whitney U-test), similar to previous observations (Wang and Movshon 2016).

### Organization of pattern motion and direction of motion within cortex

The modular organization of pattern and component neurons, together with evidence that pattern neurons integrate inputs across a broad range of component motion directions, raises the question of how these modules might be organized with respect to direction of motion columns. When examining the progression of tuning across recording depth, we noticed that while changes in tuning near reversals appeared abrupt for gratings, they were less abrupt for plaids (**Fig. 6a**). This was due to the fact that neurons near the reversals tended to have two peaks in their responses to plaids, i.e., neurons near reversals tended to be component neurons (**Fig. 6b; Fig. S5**). To assess this across recordings, we computed the ratio of component-to-pattern neurons as a function of distance from reversals. Across all sessions, we found that there were twice as many component neurons as pattern neurons near reversals (0-150 µm) whereas that ratio was reversed at sites more distant from reversals (300-450 µm) (**Fig. 6c**). To further quantify this, we fit a hyperbolic tangent function to the relationship. The amplitude of this fit (0.97) was significantly greater than that of a label-shuffled null model (permutation test, p < 0.001), and the fit crossed a 1:1 component-to-pattern ratio at a distance of 170 µm from a reversal (R^2^=0.98). Thus, component and pattern modules are not arranged randomly with respect to direction of motion columns. Instead, they are arranged systematically with respect to direction columns; component modules straddle reversals whereas pattern modules avoid them (**Fig. 6d**). This observation may account for the above evidence that pattern and component neurons represent distinct populations organized in a hierarchical circuit.

**Figure 6.**
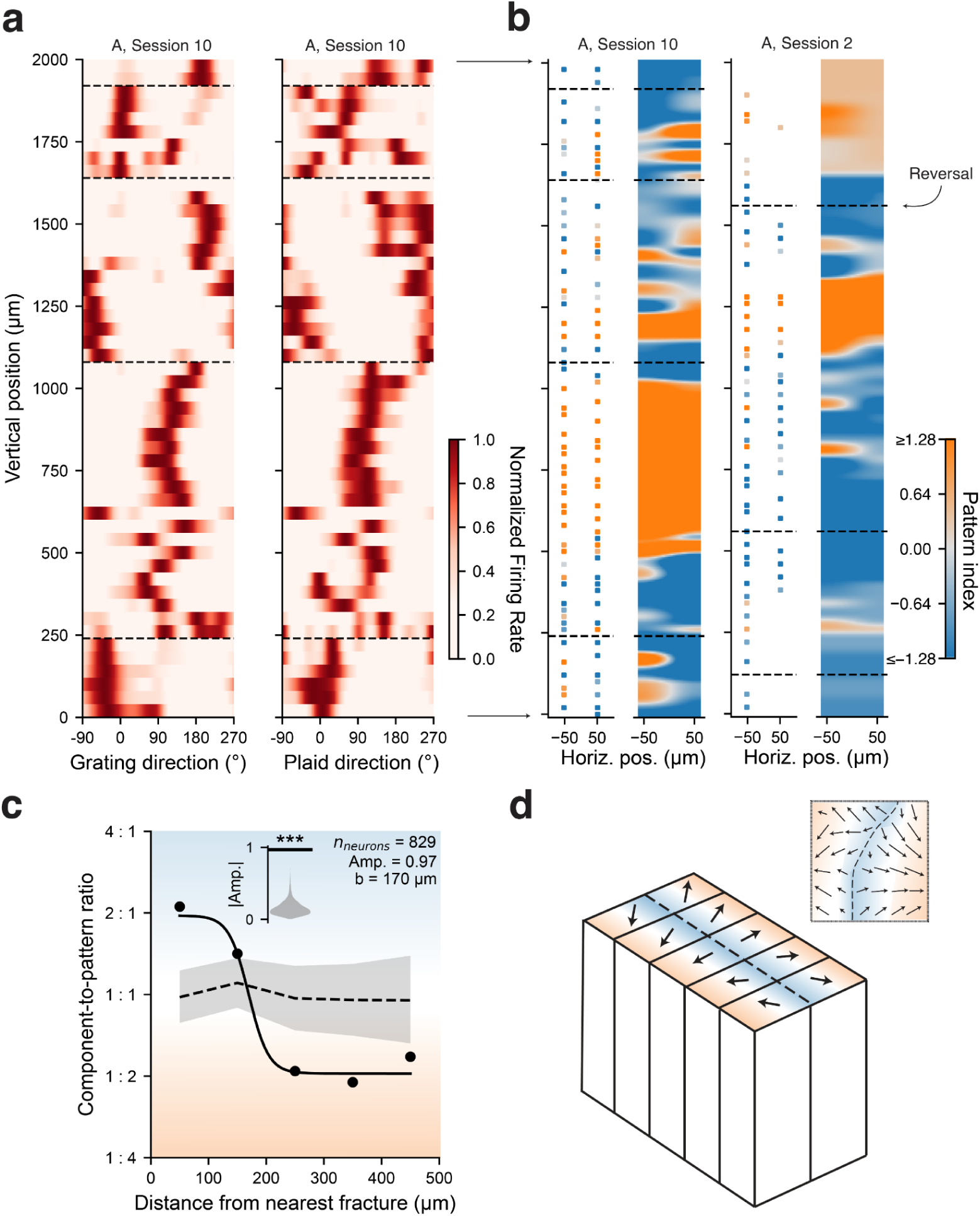
Organization of component and pattern neurons with respect to the cortical map for motion direction. **(a)** Reversals (dashed black lines) overlaid on the map of grating (left) and plaid (right) direction tuning curves for all neurons at positions relative to the probe tip in 40 μm bins in an example session. Conventions follow Fig. 1b. **(b)** Reversals overlaid on the map of pattern index for all neurons at positions relative to the probe tip in two example sessions. Conventions follow Fig. 3b. **(c)** The ratio of component to pattern neurons at different distances from reversals, aggregated across all sessions. The black solid line illustrates a tanh fit with amplitude (Amp.) and offset (b) denoted in the upper right. The black dashed line and gray shaded region denote the baseline and 95% confidence interval from a label-shuffled null model. Blue and orange shading denote regions of the y-axis where component and pattern neurons are overrepresented, respectively. The inset denotes the observed amplitude (black solid line) vs the amplitudes from the shuffled null model (*** indicate p < 0.001, permutation test). **(d)** Schematic of the organization of pattern and component neurons with respect to reversals, overlaid on models of the hypercolumnar organization of direction selectivity in area MT. Regions where component and pattern neurons are overrepresented are indicated in blue and orange, respectively.

## Discussion

Computing the direction of moving objects is a critical visual function that requires integrating the motion of component edges into a coherent pattern. Using high-density recordings in area MT, we observed a modular, hierarchical circuit for computing pattern motion. Specifically, we found that pattern and component neurons were organized into spatially distinct modules. Second, we observed that pattern and component populations could be distinguished on the basis of their spike waveforms. Third, we found that the direction of functional connections between component and pattern neurons was biased, with component neurons tending to lead pattern neurons. Fourth, pattern neurons received functional input from component neurons with a broad range of preferred directions. Lastly, we found that component and pattern modules were arranged systematically with respect to direction columns, with component modules straddling reversals and pattern modules avoiding them. Taken together, these results indicate that the representation of pattern motion emerges from a modular, hierarchical circuit composed of distinct populations of neurons organized with respect to the representation of motion direction.

### Linking cortical structure to the computation of pattern motion

In an attempt to unify our results, we propose a framework that relates the observed architecture and connectivity of pattern and component neurons to a classic solution to the problem of computing object motion. When an object moves, its pattern motion direction is consistent with a range of component motion directions within 90° of its pattern motion direction (blue vectors in **Fig. 7**) and inconsistent with component motion directions beyond 90° (red vectors in **Fig. 7**) (Marr and Ullman 1981). Each component motion signal constrains the pattern motion direction, and the intersection of these constraints uniquely determines the object’s motion (Marr and Ullman 1981; Adelson and Movshon 1982). Thus, computing object motion requires integrating inputs over a specific range of component motion directions.

**Figure 7.**
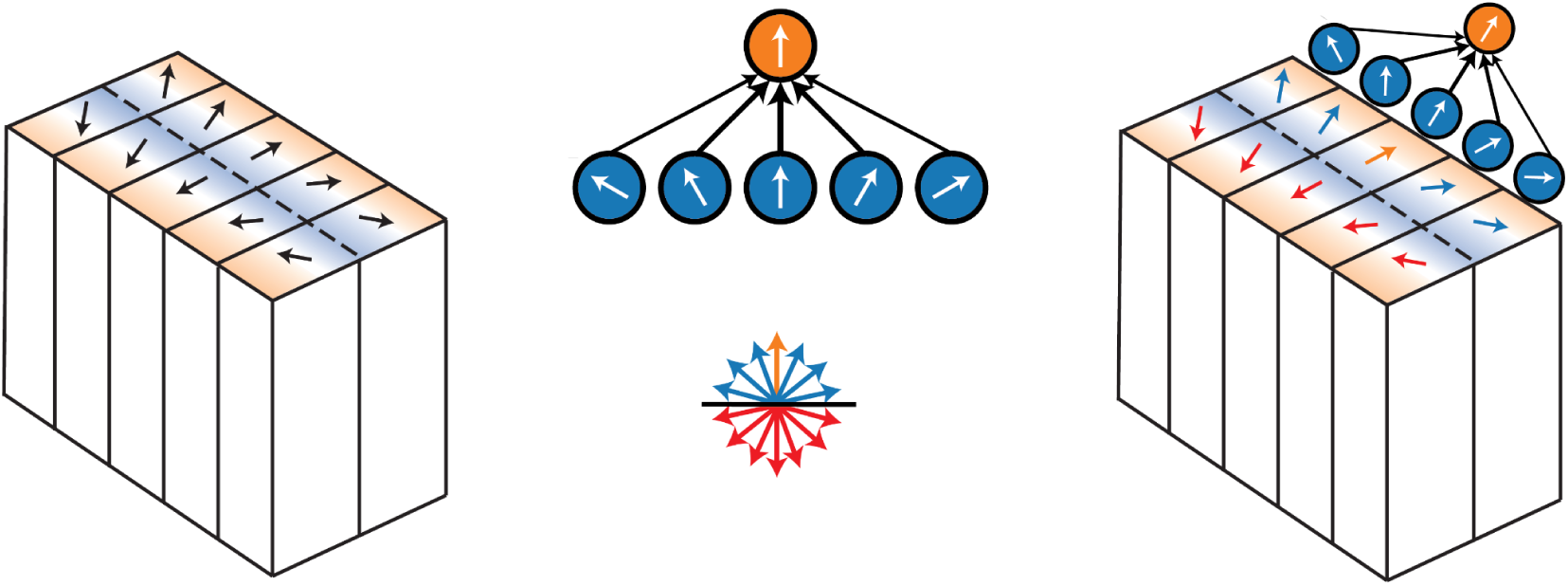
Linking cortical structure to the computation of pattern motion. Left, schematic of axis of motion columns with regions biased towards component and pattern neurons highlighted in blue and orange, respectively. Middle, top, schematic of a pattern neuron that integrates input over a range of component neurons consistent with its preferred pattern direction. Middle, bottom, schematic of the set of component directions consistent (blue) and inconsistent (red) with a particular pattern motion direction (orange) under the intersections of constraints model. Right, schematic of the predicted relationship between spatial organization and connectivity in area MT based on the intersections of constraints model. A pattern neuron (orange) positioned on one side of a reversal integrates inputs from component neurons (blue) whose preferred directions are consistent with its pattern direction (i.e., within 90°) while avoiding component neurons located across the reversal whose preferred directions are inconsistent with its pattern direction (red).

Specifically, a pattern neuron should integrate inputs from component neurons with preferred directions within 90° of its preferred pattern direction while avoiding inputs from component neurons with preferred directions outside this range. This connectivity motif predicts where pattern neurons should be situated within the columnar map for motion direction. Given that neurons tend to preferentially integrate input from their most proximal neighbors, it may be advantageous for pattern neurons to be distant from reversals. This is because the component neurons situated on the same side of the reversal as the pattern neuron are consistent with the pattern neuron’s preferred motion direction (blue vectors) while component neurons on the opposite side of the reversal have incompatible preferred directions (red vectors). Indeed, this is the architecture we observed.

### Are the observed modules laminar, columnar, or both?

Our results provide clear evidence for the presence of component and pattern modules in macaque area MT. Naturally, this raises the question of the arrangement of these modules with respect to the cortical surface. There is strong evidence that direction of motion is organized in vertical columns and that reversals in preferred direction occur when moving parallel to the cortical surface (Albright et al. 1984; Diogo et al. 2003). Given this evidence, the observed overrepresentation of component modules near reversals and pattern modules more distant from reversals suggests that there is a substantial tangential component to the component-pattern modularity. Moreover, our measurements of the rate of change in preferred direction over cortical distance are consistent with previous data from tangential penetrations (DeAngelis and Newsome 1999). Thus, our data demonstrate a substantial tangential component to the observed modularity, which aligns with earlier hints that functional cell types could be tangentially distributed in primate MT (Malonek et al. 1994; Kaskan et al. 2010; Diogo et al. 2020). Nevertheless, we emphasize that evidence for a tangential organization of component-pattern modularity is not mutually exclusive with some level of laminar organization.

Indeed, the observed differences in extracellular waveforms between pattern and component classes could be taken as evidence of a laminar organization, given that cell types are often differentially distributed across cortical layers (Callaway 1998). However, cell type can also vary tangentially to the cortical surface, often in association with functional modules. For example, in primary somatosensory cortex, barrels and intervening septa differ in their distributions of SST+ and VIP+ interneurons (Argunşah et al. 2025), and in medial entorhinal cortex, calbindin+ neurons cluster in patches tangential to the cortical surface (Ray and Brecht 2016). In primary visual cortex, morphological cell types also appear to be differentially distributed with respect to cytochrome oxidase blobs (e.g., (Rockland 2021; Nassi and Callaway 2007; Yarch et al. 2019)). Finally, in area MT, there is evidence of enhanced inhibition near reversals, which may relate to the differences between fast-spiking (putative inhibitory) pattern and component neurons observed here (Diogo et al. 2003).

### The emergence of pattern motion representations

The circuit basis for the emergence of pattern representations in area MT has been unresolved since their discovery (Wang and Movshon 2016; Rust et al. 2006; McDonald et al. 2014; Wang 1997; Movshon et al. 1985; Simoncelli and Heeger 1998; Quaia et al. 2022; Albright 1984; Rodman and Albright 1989; Diogo et al. 2020; Pack et al. 2001; Born and Bradley 2005). Some clues from past studies using single-electrode recordings are consistent with the present results. For example, pattern neurons tend to exhibit slightly longer latencies in visually evoked activity, and the onset of pattern motion signals occurs significantly later than component direction selectivity (Smith et al. 2005). However, as the authors point out, differences in latencies are consistent with a variety of possible underlying mechanisms, including differences in the local circuitry of component and pattern neurons. Latency differences could reflect either serial or parallel processing, e.g., latency differences between dorsal and ventral stream areas are thought to largely reflect parallel processes (Schmolesky et al. 1998).

Presumably, understanding the emergence of pattern representations at the circuit level requires identifying the inputs to pattern neurons. For example, knowledge of the feedforward, simple-to-complex cell hierarchy in V1 derives from evidence that complex cells receive input from simple cells (Alonso and Martinez 1998). Our results demonstrate that the direction of functional connections between component and pattern neurons is biased, suggesting component neurons provide feedforward input to pattern neurons. However, it remains possible that this circuit is not strictly serial. For example, pattern neurons in MT may also receive input from component neurons in V1 (Rodman and Albright 1989; Rust et al. 2006) or feedback from MST where pattern neurons are more prominent (Khawaja et al. 2013), and their pattern selectivity may be driven by those inputs. Nonetheless, our results substantially constrain the set of possible circuit-level models of the transformation from component to pattern representations. Specifically, models of this transformation must include a motif in which representations of pattern motion emerge from spatially distinct component and pattern modules arranged in a feedforward hierarchy.

## Methods

### Subjects

Two adult male rhesus monkeys (*Macaca mulatta*) participated in this study (Monkey A: 14 years, 11 kg, Monkey H: 13 years, 17 kg). All surgical and experimental procedures were approved by the Stanford University Institutional Animal Care and Use Committee and were in accordance with the policies and procedures of the National Institutes of Health.

### Surgical procedures and electrophysiological recordings

Monkeys were implanted with a recording chamber positioned over the superior temporal sulcus in the left (A) or right hemisphere (H). A craniotomy was performed within the chamber to provide access to the visual areas of interest. The chamber implantation and craniotomy were performed under general anesthesia and postoperative analgesics were provided during recovery.

Acute recordings in area MT were conducted using the Neuropixels 1.0 NHP probe, a high-density recording electrode optimized for use in non-human primates (Trautmann et al. 2025). In each recording session, a guide tube was first lowered to pierce the dura, then the probe was lowered through the guide tube to reach the target location. Probe position was controlled with a hydraulic (Narishige group; H) or mechanical (NAN instruments; A) microdrive system. Recordings were conducted from 384 contiguous channels along the probe to sample activity in 3.84 mm segments of tissue. The location of area MT was identified using an anatomical MRI (**Fig. S1**) and confirmed based on the physiological properties of the recorded neurons. In each of the 17 recordings (A: 13, H: 4) reported here, we observed a robust modular organization of direction selectivity along the probe shank, a hallmark of area MT (Albright et al. 1984). Given our approach to access MT and recording depth, this modular organization was most pronounced within 0-2000 µm of the probe tip across sessions in both monkeys. Subsequent analyses were therefore restricted to neurons within this depth range.

### Behavioral task and visual stimuli

Visual stimuli were presented on a ViewPixx LCD monitor (1920 x 1080 resolution, 100 Hz refresh rate) positioned 42 cm from the monkey. Stimulus presentation was controlled using the Psychophysics Toolbox (Brainard 1997) and MATLAB (MathWorks, version R2020b). Eye position was monitored at 1 kHz using the Eyelink 1000 Plus (SR Research Ltd.) eye tracker. The eye tracker was calibrated with a five-point protocol at the beginning of each session. In all tasks, monkeys initiated a trial by fixating a central dot on a gray background (33 cd/m^2^). After a prestimulus interval of 0.3-1.2 s (depending on the task and session), a sequence of visual stimuli was presented. Monkeys received a juice reward at the end of a trial for maintaining fixation throughout the trial.

*Receptive field mapping task.* In each trial of the receptive field mapping task, drifting gratings (6-13 depending on the session) were shown in a sequence, each for 0.1 s with no inter-stimulus interval. The location of each stimulus was drawn randomly from a grid with 3° spacing, covering vertical positions from-15° to 15° and horizontal positions from-3° to 15° for monkey A (right visual field), and from-21° to 0° for monkey H (left visual field). The drift direction of the stimulus was drawn randomly from one of eight directions in 45° increments spanning 0-315° (drift speed 15°/sec). The gratings (1 cycle/°) were modulated with a Gaussian envelope (sigma 1°). Receptive fields were mapped online with threshold crossing to guide stimulus positioning and were analyzed further offline.

*Pattern motion task.* In each trial of the pattern motion task, three drifting grating and plaid stimuli were shown in a sequence, each for 0.25 s with an inter-stimulus interval of 0.15 s. Stimuli consisted of drifting gratings and plaids constructed by superimposing two such gratings with drift directions separated by 120°. Stimulus drift direction and type (grating vs plaid) were randomly selected on each presentation. Stimuli drifted in one of twelve directions in 30° increments spanning 0-330° (drift speed 10°/sec; 1 cycle/deg). Stimulus size and location were selected in each session to ensure coverage of the population receptive field. In 6 sessions (A: 5, H: 1), stimuli consisted of gratings (50% Michelson contrast) presented inside a circular aperture, and plaids constructed by superimposing two such gratings with equal weights on the two components, following previous studies (Smith et al. 2005) (**Fig. S6a**). In 11 additional sessions (A:8, H:3), stimuli consisted of gratings (33% Michelson contrast) presented inside a Gaussian envelope, and plaids constructed by superimposing two such gratings with relative weights of the two components derived from a Gaussian (**Fig. S6b**). Results were consistent across the two plaid stimulus sets and thus were pooled for all analyses (**Fig. S7**). In some sessions, four additional triplaid stimuli (Jazayeri et al. 2012) were interleaved with gratings and plaids, but are not analyzed further here.

### Neural data preprocessing

Action potential band data was spike-sorted using Kilosort4 with default parameters to obtain spike times for individual neurons and waveform templates (Pachitariu et al. 2024). All single and multi-units identified by Kilosort4 were aggregated and are referred to as ‘neurons’ hereafter. Neurons were included in subsequent analyses if they had a waveform consistent with a somatic spike and exhibited reliable, direction-selective visual responses. A waveform was considered somatic if it had a positive trough-to-peak duration with a larger trough than peak amplitude. A neuron was considered visually-responsive and direction-selective if its firing rate in its preferred drifting grating condition exceeded 5 Hz, the baseline rate (one-sided Mann–Whitney U test, p < 0.05), and the rate evoked by the antipreferred drifting grating (direction selectivity index ≥ 0.25). The direction selectivity index (DSI) is:

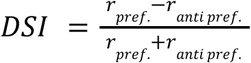

where 𝑟 _𝑝𝑟𝑒𝑓_ is the neuron’s firing rate to its preferred grating direction and 𝑟 _𝑎𝑛𝑡𝑖 𝑝𝑟𝑒𝑓_ is its firing rate to the opposite grating direction. This yielded a total of 2,096 neurons across all sessions, with a mean of 123 neurons per session (min = 53, max = 184).

### Classification of component and pattern neurons

Neurons were classified as component or pattern neurons following previous studies (Smith et al. 2005; Wang and Movshon 2016; Movshon et al. 1985). Specifically, we computed grating and plaid direction tuning curves by averaging responses in each stimulus condition across trials and time (30-250 ms after stimulus onset). We then computed the partial correlation between the measured plaid tuning curve and predictions of the pattern and component models. The pattern model prediction is the tuning curve for gratings whereas the component model prediction is the sum of the tuning curve for gratings rotated by 60° and-60°. The partial correlations for the pattern and component predictions are:

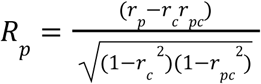

and

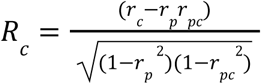

where 𝑟*_p_* and 𝑟*_c_* are the correlations of the measured plaid tuning curve with the pattern and component predictions, respectively, and 𝑟*_pc_*is the correlation of the two predictions. Finally, the partial correlations were converted into Z-scores using the Fisher z-transformation:

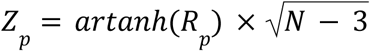

and

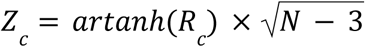

where 𝑎𝑟𝑡𝑎𝑛ℎ is the inverse hyperbolic tangent function and 𝑁 is the number of directions in the tuning curve (here, 𝑁=12). The pattern index is given by 𝑃𝐼 = 𝑍*_p_* − 𝑍*_c_*. Neurons with 𝑃𝐼 ≥ 1. 28 and 𝑍*_p_* ≥ 1. 28 were classified as pattern neurons, and neurons with 𝑃𝐼 ≤− 1. 28 and 𝑍*_c_* ≥ 1. 28 were classified as component neurons following criteria used in previous studies (Smith et al. 2005). Neurons that did not meet these criteria were labeled unclassified (Wang and Movshon 2016; Smith et al. 2005).

### Functional connectivity analyses

To analyze functional connectivity, we computed the cross-correlation between all pairs of simultaneously recorded neurons (English et al. 2017; Stark and Abeles 2009). We focused on activity in the late stimulus window (100-250 ms after stimulus onset) to ensure the analysis was not affected by the visual transient. The cross-correlation histogram (CCH) for a pair of neurons (𝑗, 𝑘) was defined as:

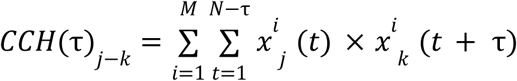

where 𝑀 is the number of trials, 𝑁 is the number of 1 ms time bins within a trial, τ is the time lag, 𝑥^𝑖^ *_j_*(𝑡) is 1 if neuron 𝑗 fired in time bin 𝑡 of trial 𝑖. The CCH is computed with neuron 𝑗 as the source neuron and 𝑘 as the target neuron, denoted 𝑗-𝑘.

To identify functionally connected pairs of neurons, we compared the measured CCH to a slow baseline CCH computed by convolving the measured CCH with a partially hollowed Gaussian kernel with a standard deviation of 10 ms and hollow fraction of 0.6 (English et al. 2017; Stark and Abeles 2009). An opportunity correction was applied to the slow baseline to account for the degree of overlap of the two spike trains for each time lag. A pair of neurons was classified as functionally connected if the peak of the CCH fell within 1-5 ms of zero time lag and was significantly larger than both the slow baseline CCH at the corresponding time lag and the CCH at time lags from-5-0 ms.

More specifically, the probability of observing a CCH value of 𝑛 or more at lag τ given the slow baseline CCH at lag τ was estimated using a Poisson distribution with a mid-p correction (English et al. 2017):

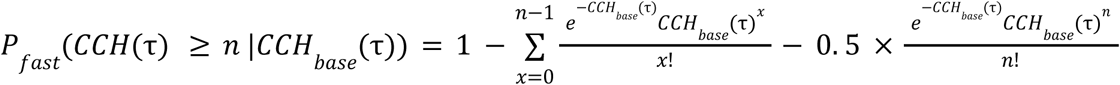

Similarly, the probability of observing a CCH value of 𝑛 or more at lag τ given the maximum value of the CCH at time lags from-5-0 ms, λ = 𝑚𝑎𝑥 _τ∈[−5, 0]_ 𝐶𝐶𝐻(τ), was estimated using a Poisson distribution with a mid-p correction (English et al. 2017):

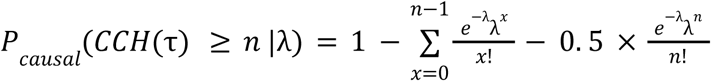

A pair of neurons was labeled as functionally connected if 𝑃_𝑓𝑎𝑠𝑡_ was less than 0.01 and 𝑃_𝑐𝑎𝑢𝑠𝑎𝑙_ was less than 0.01, following criteria used in previous work (English et al. 2017). Results are robust to the inclusion of the early visual period (**Fig. S4**). We note that although the criteria used here for identifying functional connections were developed specifically to identify monosynaptic connections (English et al. 2017), functional connectivity results may not always reflect synaptic connectivity and should be interpreted with caution.

### Reversal detection

We utilized an automated approach to identify discontinuities (‘reversals’) in the cortical map for preferred direction. Neurons were binned in 40 µm increments along the probe depth, and each bin was assigned the preferred direction of its highest firing rate neuron. Locations where the difference in preferred direction between adjacent bins exceeded 100° were selected as candidate reversals. These candidates were then refined by requiring local stability on each side of the reversal (adjacent bins where difference in preferred direction was less than 50°) and local instability across the reversal (difference in median preferred direction of 3-bin windows on each side of the reversal exceeded 100°). 14/17 sessions were included for subsequent reversal analyses; the remaining sessions exhibited substantial scatter in preferred direction that precluded reliable reversal estimation. Reversals automatically detected using this approach are illustrated for example sessions in **Fig. 1** and **Fig. S2**.

### Waveform preprocessing, feature extraction, and classification

Waveform analyses were performed on the subset of cells classified as component or pattern neurons. Prior to classification, waveforms were baseline subtracted, normalized by their trough amplitude, and aligned to the trough time (“preprocessed waveform”). The input to the classifier was the preprocessed waveform, where the number of input features is the number of samples in the waveform (here, 51). The output of the classifier was a binary label corresponding to pattern or component. The classifier was a support vector machine with a radial basis function kernel (scikit-learn, (Pedregosa et al. 2012)) which can learn flexible nonlinear mappings between waveform shape and functional class. The classifier was trained and tested with 5-fold cross-validation.

Nested cross-validation was used to select the regularization hyperparameter for the classifier on each fold. Class-balanced accuracy is reported in all figures (scikit-learn, (Brodersen et al. 2010; Pedregosa et al. 2012)). To assess statistical significance, observed accuracy was compared to a null model where component and pattern labels were shuffled. Trough-to-peak duration and normalized peak amplitude features were extracted from preprocessed waveforms. The normalized peak amplitude is defined as the maximum of the preprocessed waveform. Waveforms with trough-to-peak durations between 0 and 200 µs were classified as fast-spiking (FS), and those with trough-to-peak durations greater than 200 µs were classified as regular-spiking (RS), following criteria used in previous studies (Mitchell et al. 2007).

### Curve fitting and statistics

To quantify the modular organization of tuning properties, we fit exponential decay functions of the form, 𝑦 = 𝑏 ± 𝐴 × 𝑒^−𝑥/τ^, where 𝑥 is the distance between a pair of neurons and 𝑦 is their difference in preferred direction (**Fig. 1c**) or a binary indicator of whether they share the same pattern/component label (**Fig. 3c**). 𝐴 corresponds to the amplitude, 𝑏 corresponds to the bias, and τ corresponds to the spatial scale of the modularity. To quantify the relationship between pattern modularity and reversals, we fit a hyperbolic tangent function of the form, 𝑦 = 𝐴 𝑡𝑎𝑛ℎ((𝑥 − 𝑏)/𝑇), to the binned data, where 𝑥 is the distance of a neuron from its nearest reversal and 𝑦 is the log ratio of component to pattern neurons (**Fig. 6c**). 𝐴 corresponds to the amplitude, 𝑇 to the slope, and 𝑏 to the distance from a reversal with a 1:1 ratio of component to pattern neurons. Fit parameters were estimated using nonlinear least squares (scipy.optimize.curve_fit; (Virtanen et al. 2020)). The statistical significance of modularity results was assessed using permutation tests with label-shuffled null models. Specifically, we permuted the preferred direction or pattern/component labels for neurons within each session, and then repeated each analysis to obtain a shuffled baseline and 95% confidence interval. We also fit the corresponding function to each shuffled curve to obtain a distribution of fit amplitudes under the null model. p-values were computed as the fraction of shuffles yielding amplitudes greater than or equal to the observed value. Nonparametric tests were used to assess asymmetries in functional connectivity (binomial test; **Fig. 5b**) and differences in waveform metrics (Mann-Whitney U-test; **Fig. 4g**).

## Data and code availability

The data and code supporting this work will be made available upon publication.

## Acknowledgements

We thank William T. Newsome, Thomas D. Albright, Kristina Nielsen, and J. Anthony Movshon for valuable feedback and suggestions on this work. This work was supported by NIH EY014924, NS116623 and a Ben Barres Professorship to T.M., and NSF GRFP 2146755 and a Stanford Bio-X Graduate Student Fellowship to E.T.

## Author Contributions

E.T. and T.M. designed research; E.T., R.X., S.Z., S.S., D.L., S.C., and T.M. performed research; E.T. and C.Y. analyzed data; and E.T. and T.M. wrote the paper.

## Supplementary Information

**Supplementary Figure 1.**
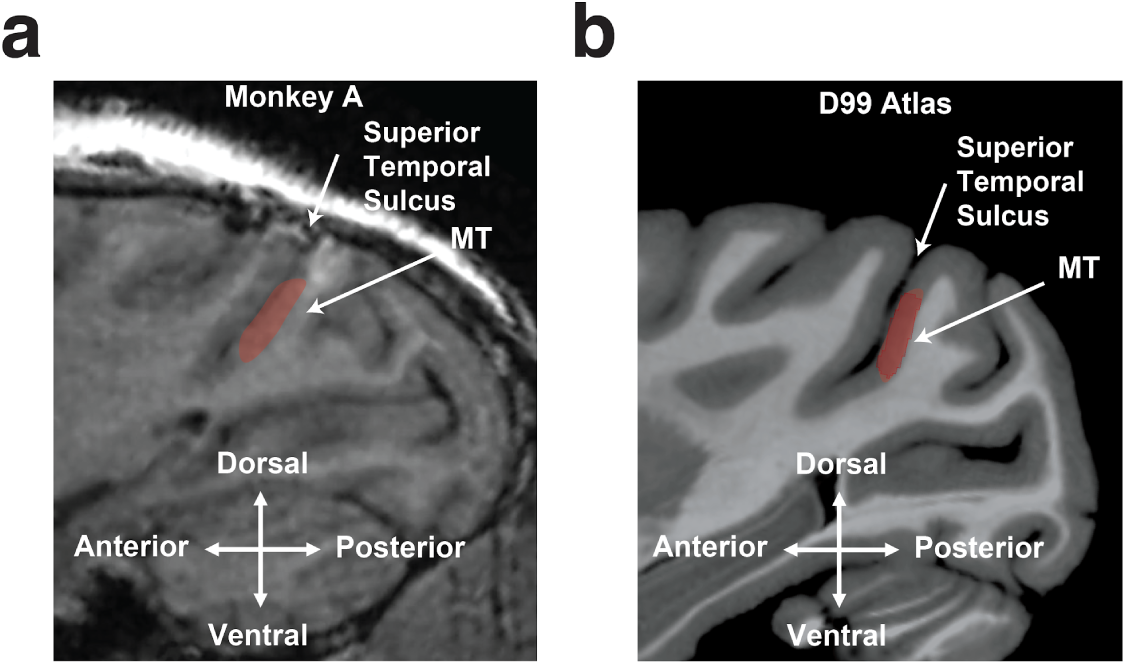
MR image of the superior temporal sulcus and area MT in monkey A and the D99 atlas. Sagittal MR image from monkey A (a) and the D99 macaque brain atlas (b). The location of the superior temporal sulcus and area MT (red shaded) are illustrated.

**Supplementary Figure 2.**
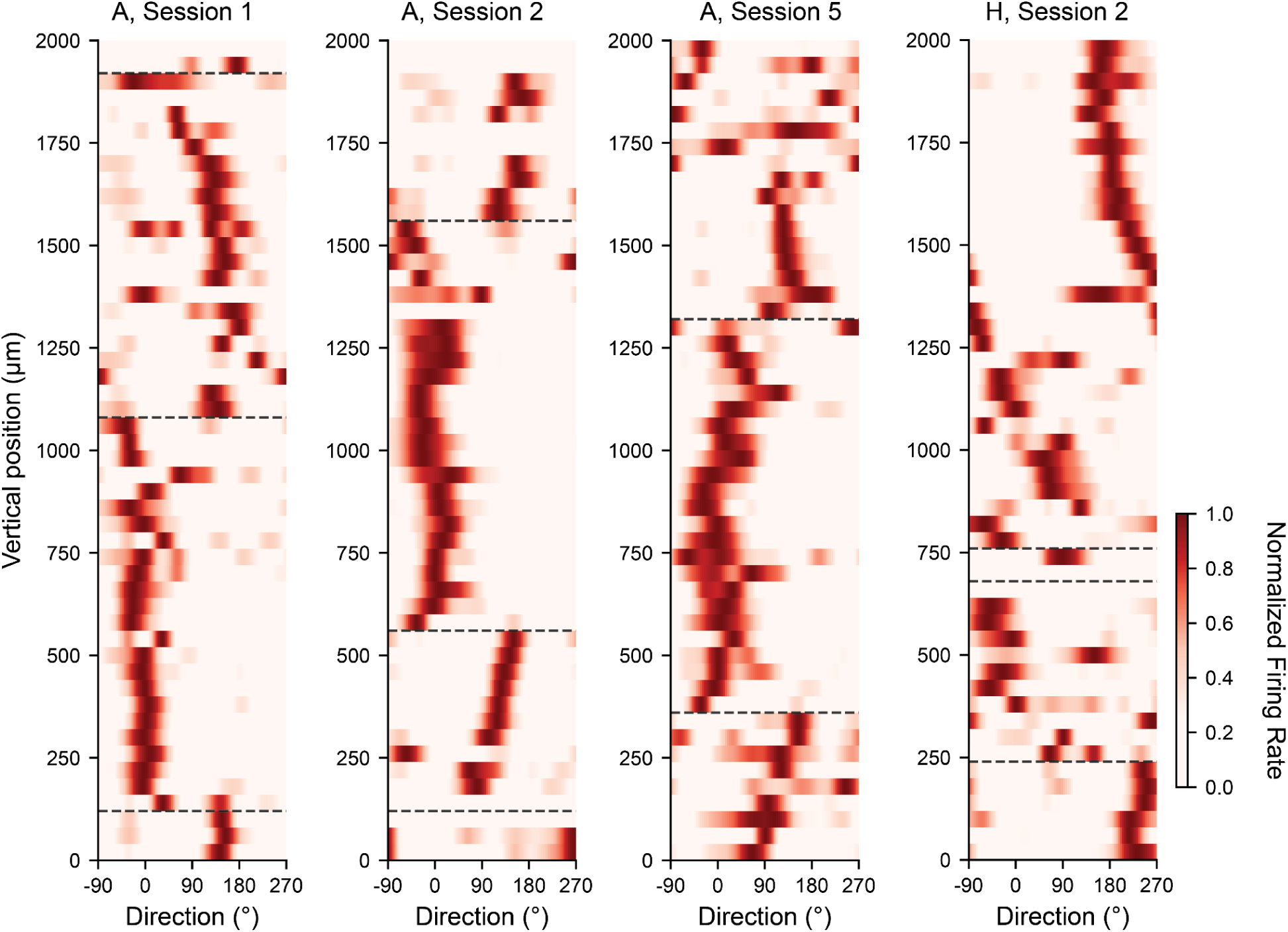
Additional examples of modular organization of direction tuning in area MT. Grating direction tuning curves for all neurons at positions relative to the probe tip in 40 μm bins in four additional example sessions. Conventions follow Fig. 1b.

**Supplementary Figure 3.**
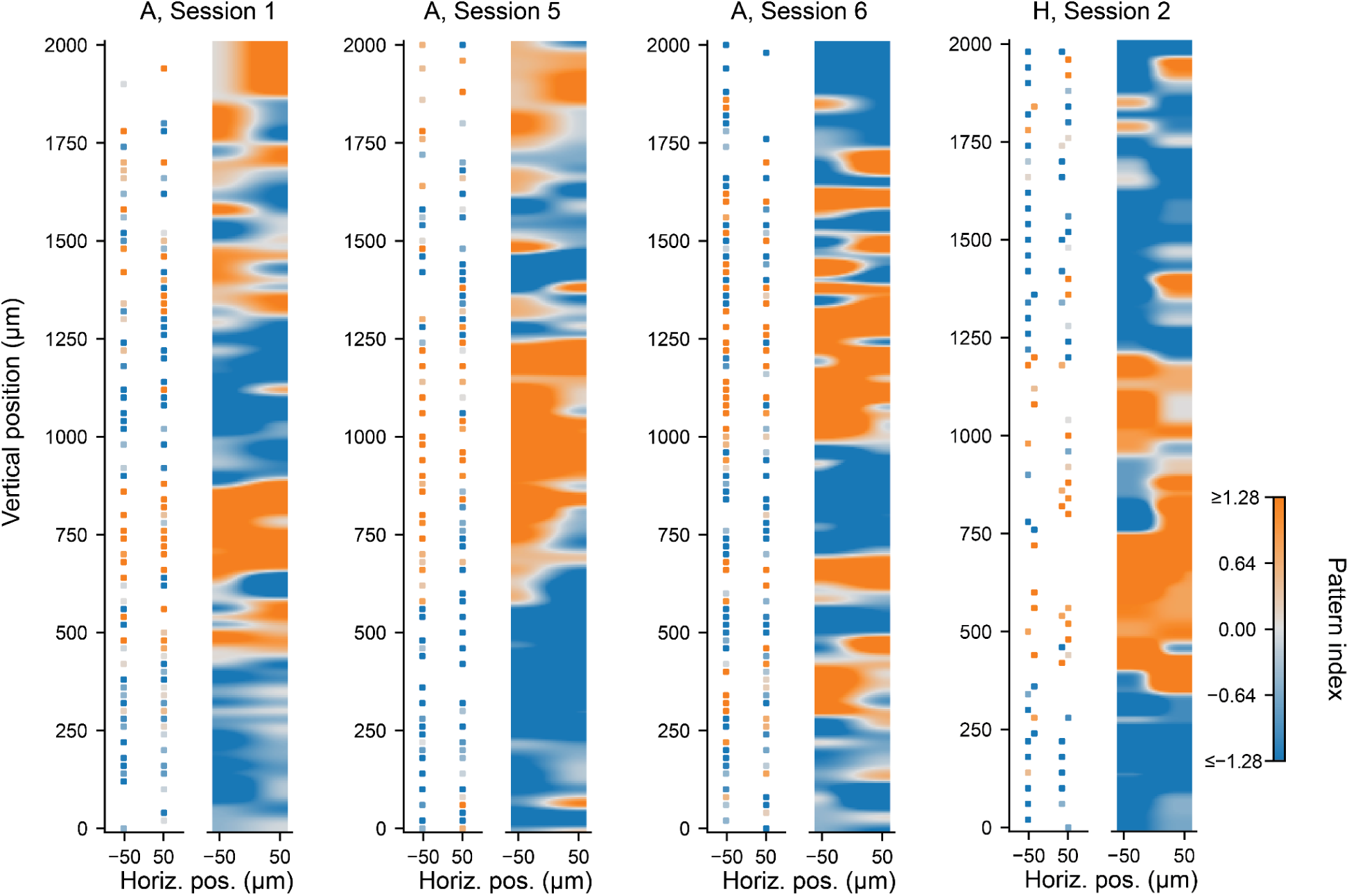
Additional examples of modular organization of pattern and component neurons. Pattern index for all neurons at positions relative to the probe tip in four additional example sessions. Conventions follow Fig. 3b.

**Supplementary Figure 4.**
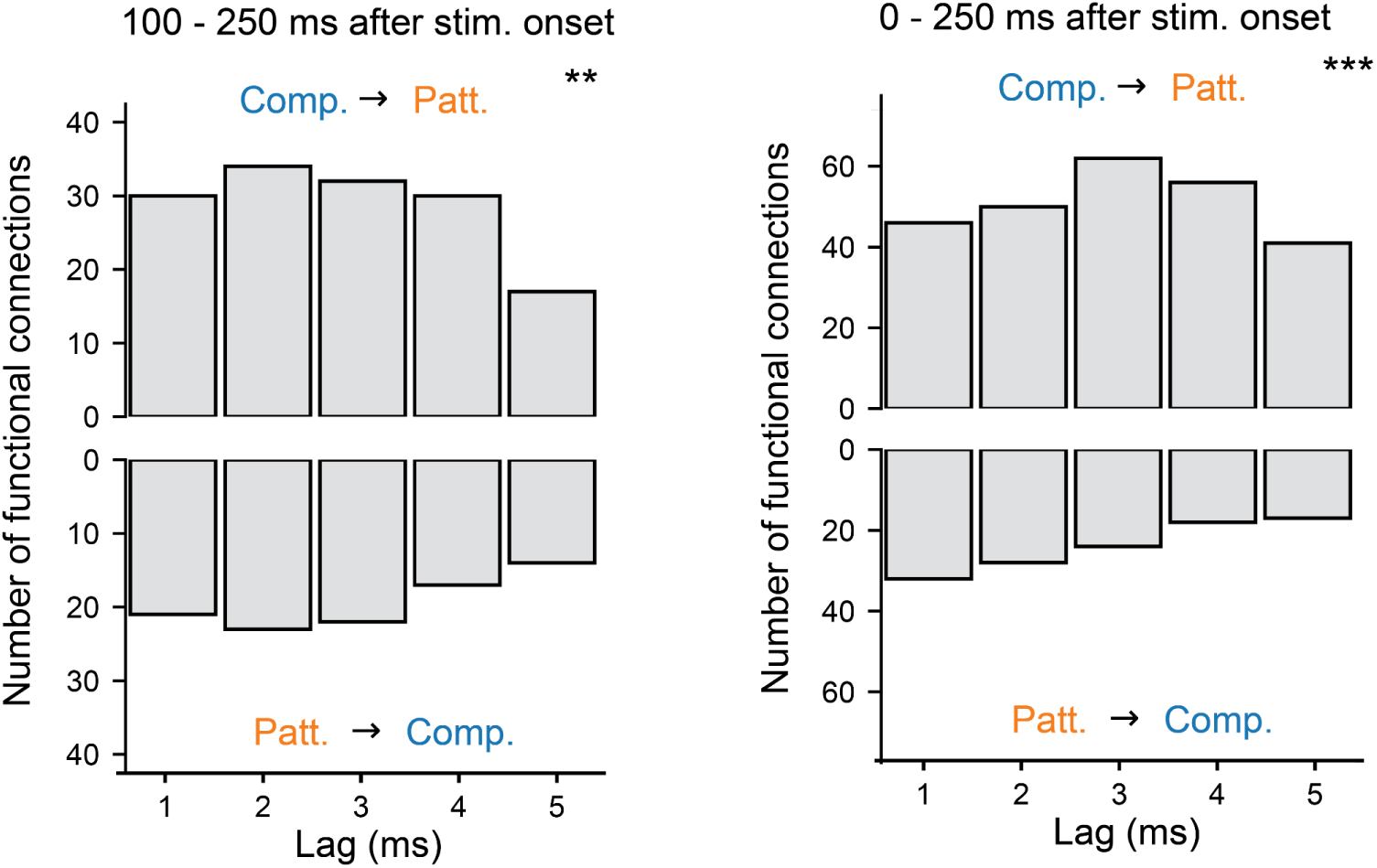
Functional connections between component and pattern neurons, computed excluding or including the early visual period. Functional connection asymmetry is robust to the inclusion of the early visual period. The distribution of lags for component-to-pattern and pattern-to-component functional connections is illustrated across all sessions. Functional connectivity is computed using the time interval from 100 ms - 250 ms after stimulus onset (left) and 0 ms - 250 ms after stimulus onset (right). Conventions follow Fig. 5b.

**Supplementary Figure 5.**
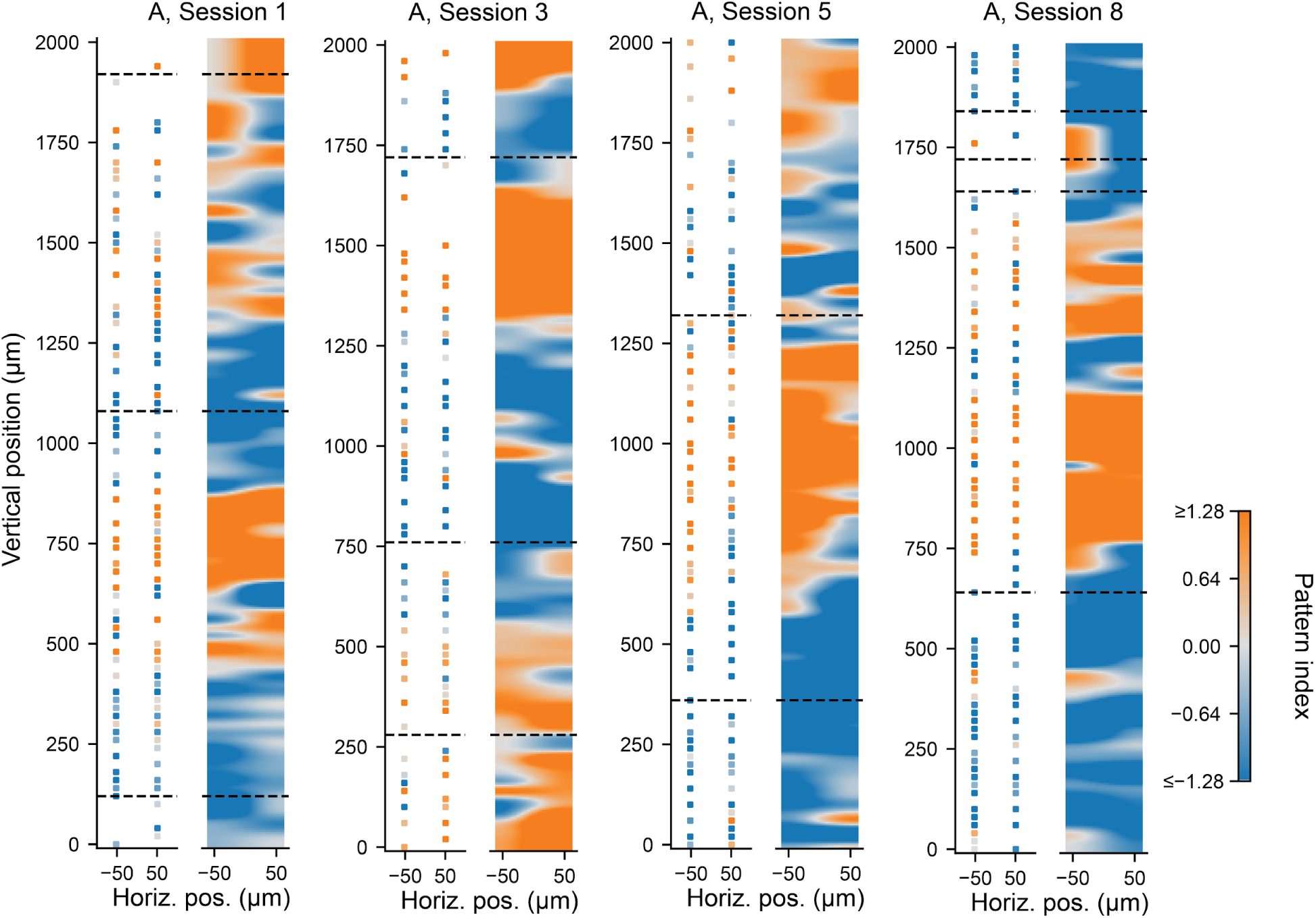
Additional examples of organization of component and pattern neurons with respect to the cortical map for motion direction. Reversals (dashed black lines) overlaid on the map of pattern index for all neurons at positions relative to the probe tip in four additional example sessions. Conventions follow Fig. 6b.

**Supplementary Figure 6.**
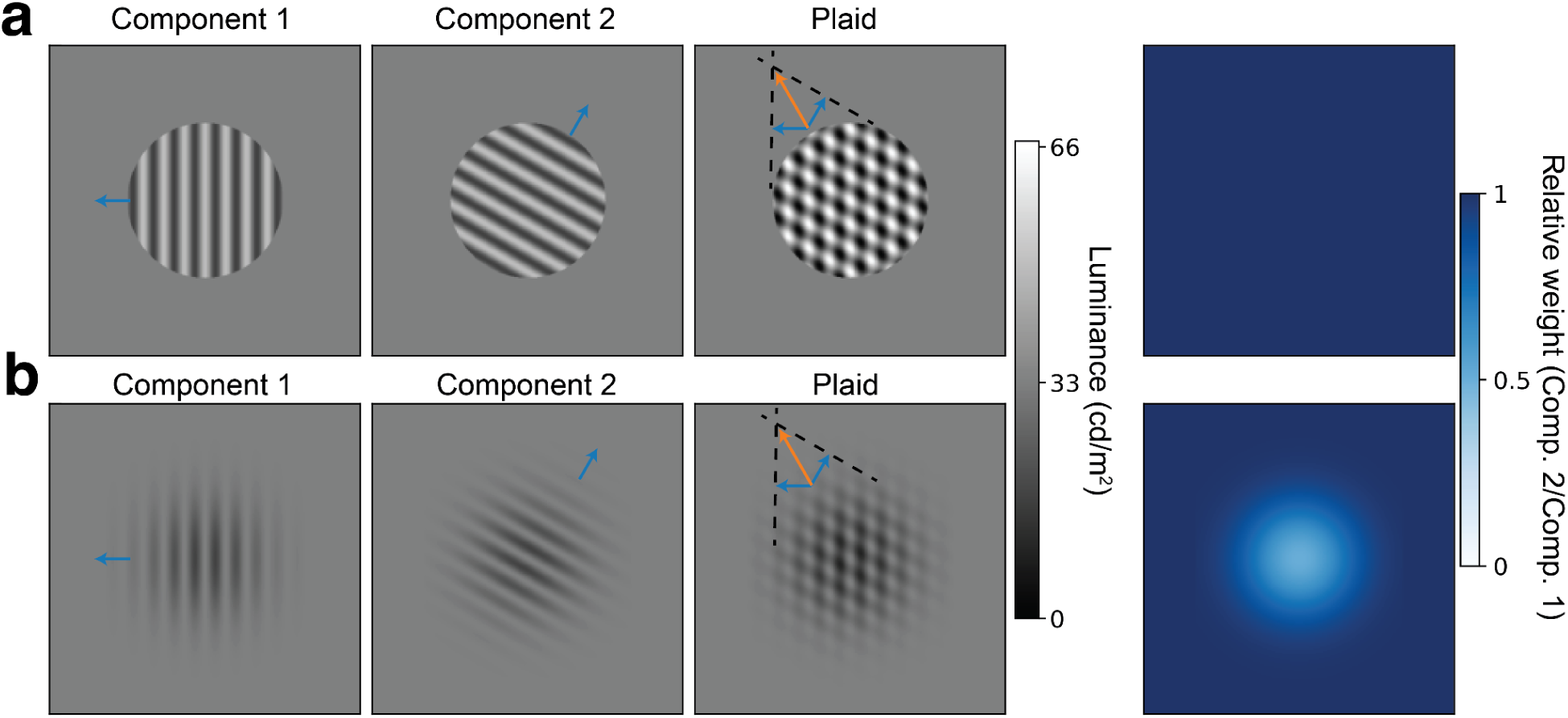
Grating and plaid stimuli for measuring pattern and component motion selectivity. (**a**) First plaid stimulus set (A: Sessions 1-5, H: Session 1). Two example component gratings, offset by 120°, and corresponding plaid stimulus constructed by superimposing the two components with equal weights. The blue arrows denote the component motion speed and drift direction. The dashed lines denote the constraints induced by component motion, and the orange arrow denotes the pattern motion vector derived from the intersection of constraints model. (**b**) Second plaid stimulus set (A: Sessions 6-13, H: Session 2-4). Two example component gratings, offset by 120°, and corresponding plaid stimulus constructed by superimposing the two components with the relative weights of the components derived from a Gaussian envelope that varies between 0.5 and 1. Conventions are similar to (a). All plaid stimuli used in this study can be described as a weighted sum of the two components, 𝑃 = (𝐶_1_ − β) + α(𝐶_2_ − β) + β, where β is the background luminance, 𝐶_1_ and 𝐶_2_ are the two components, and α is the relative weight of 𝐶_2_ to 𝐶_1_. In the first stimulus set (a), α is set to unity, and in the second stimulus set (b), α is determined by a Gaussian envelope.

**Supplementary Figure 7.**
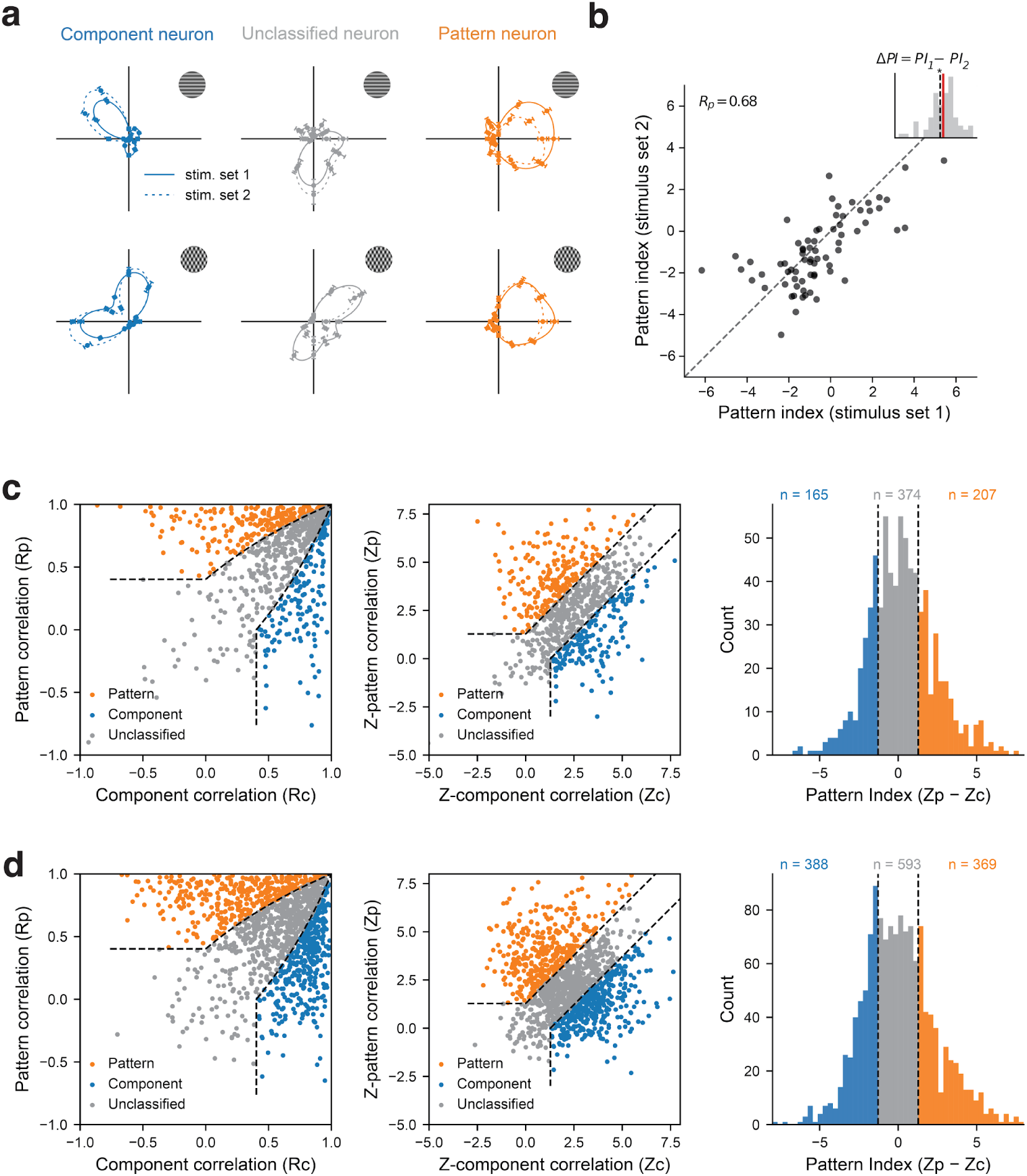
Tuning to grating and plaid stimuli and classification of pattern and component neurons, split by stimulus set. (**a**) Tuning curves to gratings (top) and plaids (bottom) for three example neurons when presented with the two sets of component and pattern motion stimuli tested in this study in the same session (see **Fig. S6**). Solid lines denote stimulus set 1 and dashed lines denote stimulus set 2. Error bars denote SEM. (**b**) Correlation between the pattern index computed using stimulus set 1 versus stimulus set 2 in the same session. The Pearson correlation coefficient and identity line (dashed black) are denoted. Inset illustrates the pairwise difference in pattern index between the two stimulus sets, with mean difference denoted by the red line and 0 denoted by the dashed black line. Asterisk denotes significant difference from 0 (Wilcoxon signed-rank test, mean diff.=0.34, p < 0.05). **(c,d)** Classification of pattern and component neurons in sessions where stimulus set 1 (c) and stimulus set 2 (d) was presented. Conventions follow Fig. 2c.

